# Organizing the coactivity structure of the hippocampus from robust to flexible memory

**DOI:** 10.1101/2023.09.26.558975

**Authors:** Giuseppe P. Gava, Laura Lefèvre, Tabitha Broadbelt, Stephen B. McHugh, Lopes-dos-Santos Vítor, Demi Brizee, Katja Hartwich, Hanna Sjoberg, Pavel V. Perestenko, Robert Toth, Andrew Sharott, David Dupret

## Abstract

New memories are integrated into prior knowledge of the world. But what if consecutive memories exert opposing demands on the host brain network? We report that acquiring a robust (food-context) memory constrains the hippocampus within a population activity space of highly correlated spike trains that prevents subsequent computation of a flexible (object-location) memory. This densely correlated firing structure developed over repeated mnemonic experience, gradually coupling neurons of the superficial CA1 *pyramidale* sublayer to whole population activity. Applying hippocampal theta-driven closed-loop optogenetic suppression to mitigate this neuronal recruitment during (food-context) memory formation relaxed the topological constraint on hippocampal coactivity and restored subsequent flexible (object-location) memory. These findings uncover an organizational principle for the peer-to-peer coactivity structure of the hippocampal cell population to successfully meet memory demands.

## Main Text

Every day, we use our existing knowledge to guide the actions we make in our environment, integrating new information to gain further knowledge about the world. Therefore, building new memories does not take place in a state of *tabula rasa*, but against a background of prior experiences that have been accumulated across the lifespan and have shaped their host brain networks [1,2].

The hippocampus network uses the collective activity of the population of its neurons to support everyday memory [3,4]. In principle, the level and structure of the activity coupling between individual neurons could reflect a critical tradeoff between the robustness versus the flexibility of the whole population in processing information. That is, strong peer-to-peer coupling could yield highly correlated spike trains, increasing the consistency of activity patterns within the population for robust memory expression. In contrast, weaker population coupling could release network activity space for new patterns, allowing more diverse mnemonic representations for dynamically adaptable behavior. However, the hippocampus may have to switch between robust versus flexible computations depending on current demands. What are the consequences of placing the hippocampal population into a robust computational mode for subsequent memories that instead require flexible information processing?

To address this question, we trained mice to first acquire a strong contextual memory. On every day of our paradigm (16 consecutive days), mice explored two arenas (Figure 1A, B). During the first 10 days (‘Food-context conditioning’), we paired one arena (context *X*) with two regular-diet (Chow) pellets (Figure S1A). The other arena (context *Y*) contained one chow pellet and one high-fat-diet (Hfd) pellet (Figure S1A), which mice encountered for the first time and did not eat much on day 1 (Figure 1C). By repeating these foraging sessions on each subsequent conditioning day, mice showed escalated food intake in context *Y* (Figure 1C; from day 1 to day 10, a fold-change of 17.47 ± 5.34 versus 0.72 ± 0.26 in context *Y* versus *X*; mean ± s.e.m.). To probe discriminative food-context association, we then measured their propensity to express context-biased feeding. By providing both arenas with new food items in post-conditioning days (‘Novel food test’; days 11 and 12; Figure 1B and Figure S1B), we observed higher novel food intake in context *Y* compared to context *X* (Figure 1D; a fold-change of 2.71 ± 1.07 in context *Y* versus *X*; mean ± s.e.m.). Thus, mice in the Hfd-conditioned context readily overcame the rodent natural tendency to express food neophobia [5,6]. Mice also exhibited lower Hfd intake when provided in a third arena never paired with any food (Figure S1C). Body weight remained stable across task days (Figure S1D). Thus, mice acquired a discriminative memory that shaped their behavior in a context-dependent manner.

**Figure 1.**
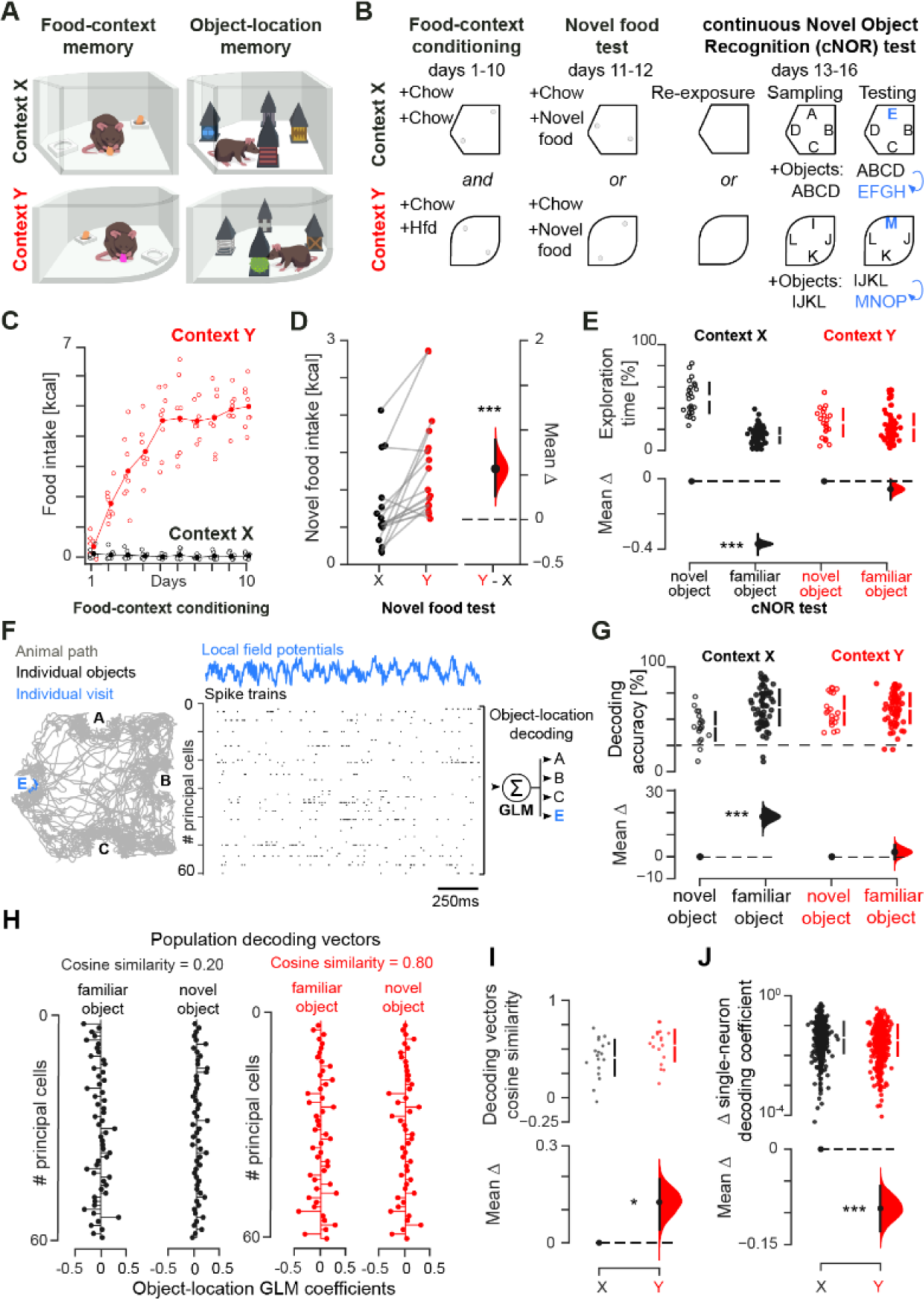
Robust contextual (food) memory prevents subsequent flexible (object) memory. **(A, B)** Behavioral tasks with open-field contexts **(A)** and multiday layout **(B)** for a two-memory paradigm. **(C)** Animals’ food intake during contextual conditioning (each data point represents one mouse). (**D)** Estimation plot (see Methods) showing the effect size for the difference in novel food intake across context *X* and *Y* after conditioning. (**E**) Likewise, shown is the percentage of exploration time with novel and familiar objects during cNOR tests in context *X* or *Y*. (**F**) Schematic of the GLM employed to predict the identity of each object-location compound from ongoing population vectors of theta-nested principal cell spiking. For clarity, a sample of a few seconds of data for one object visit (blue trace) is shown. (**G**) Estimation plot showing the classification accuracy of object-location compound in test *n* by GLM trained in session *n* − 1 (each data point represents one object). **(H)** Example population decoding vector pairs containing the neuron-wise GLM coefficients for the familiar versus a novel object at the same location in either context *X* or *Y* (each data point represents one neuron). **(I, J)** Estimation plots showing the cosine similarity between familiar and novel object-location GLM vectors **(I)** and the change (update) in single neuron contribution to whole-population novel object-location classification **(J)** across two consecutive cNOR tests in either context. ***P<0.001, *P<0.05

Having instantiated this discriminative contextual memory, we next switched task demands to assess novelty detection in these contexts previously paired with food. For this, we used a hippocampus-dependent continuous Novel Object Recognition task (‘cNOR test’; from day 13 to day 16; Figure 1A, B and Figure S2A-D). On each cNOR day, we re-exposed mice to either context *X* or *Y* (‘Re-exposure’; without any food) before they encountered four novel objects there (‘Sampling’; Figure 1A, B and Figure S2A, B). Across four more exploration sessions that day, we iteratively replaced one of the initially sampled objects with a new one (‘Testing’). This procedure thus yielded a set of three familiar (already-seen) objects and one novel (first-time-seen) object in each cNOR test (Figure 1B and Figure S2A). We measured novelty detection in cNOR test *n* using the time spent exploring the novel object over the total time spent on all four objects, thereby probing memory for objects explored in session *n* − 1. Mice showed novel object preference in context *X* but not in context *Y* (Figure 1E). Locomotor speed and distance travelled did not differ across contexts (Figure S2E-G). Thus, the robust (food) memory acquired in context *Y* interfered with a subsequent flexible (object-location) memory in that context.

We thus aimed to identify the neuronal correlates of this cross-memory interference. During active behavior, groups of principal cells recruited from the population of hippocampal neurons cooperate within the timeframe of theta-band (5–12Hz) oscillations to support codes and computations for memory [3]. We recorded cell ensembles and local field potentials in the CA1 *stratum pyramidale* of these mice. Using the action potentials discharged by principal cells in theta cycles during exploration of each object in cNOR days (Figure 1F and Figure S2H), a generalized linear model (GLM) trained on session *n* − 1 and applied in test *n* identified each object-location compound with up to 93.5% accuracy (range 9.4 – 93.5 %; mean 55.0 % compared to a chance level of 25.0 %; mean ± s.e.m. number of principal cells = 51.1 ± 3.7 per GLM), consistent with work showing population-level object representation in the hippocampus [7]. In context *X*, the mean accuracy of this population-level classification started at 38.6 ± 3.2 % for the novel object-location compound and improved as the object gained familiarity in subsequent tests (Figure 1G). This across-test gain in object-location representation did not occur in context *Y*. In fact, the mean classification accuracy there started at higher levels for the novel object-location (54.9 ± 3.4 %; P<0.001, permutation test, compared to context *X*), without significant changes in the following tests (Figure 1G). Compared to context *X*, the contribution of individual cells to each novel object-location classification during test *n* in context *Y* resembled its previous one expressed in session *n* − 1 at that location while having another (familiar) object. This was reported by the higher similarity between the population decoding vector that contained the set of neuron-wise GLM coefficients representing the novel object during test *n* in context *Y* versus that representing the familiar object at the same location during session *n* − 1 (Figure 1H, I). In line with this observation, the across-test modulation in single-neuron contributions to population decoding vectors when encountering a novel object (i.e., the changes in the magnitude of individual GLM coefficients) was weaker in context *Y* compared to *X* (Figure 1J). These results suggested a population representational rigidity prevented the hippocampus from updating object-location memory in context *Y*.

We hypothesized that this acquired representational rigidity reflects the organization of the population activity into a non-permissive structure. That is, Hfd context-conditioning yielded highly correlated firing patterns that created a dense network activity space for strong contextual (food) memory. But this later conflicted with the switch to a different demand where continually processing familiar versus novel stimuli would instead require sparser, weakly correlated patterns for disentangling discrete (object-location) representations. We indeed found that the correlational structure of the population activity was markedly different in context *Y* compared to *X* (Figure 2). We quantified the coactivity association of each cell pair (*i*, *j*) by predicting the theta-nested spike discharge of neuron *j* with the activity of neuron *i* while regressing out the activity of the remaining population (Figure 2A). This procedure returned a matrix of β regression weights (Figure 2B) that represented the neurons pairwise coactivity structure of the population in each context. With this, for both context *X* and *Y* we constructed weighted neuronal graphs (with no self-connections) where each node is a cell and the edge linking any two nodes represents the coactivity of that cell pair (Figure 2B, C; n = 108 graphs and 210,984 coactivity pairs). We found that neuronal graphs contained more triads of coactive nodes in context *Y* than *X*, as shown by higher clustering coefficients (Figure 2D, E). Moreover, the population coactivity strength level, calculated for each node as the average weight of all its edges, was also higher in *Y* (Figure S3A), despite no difference in the mean neurons’ firing rate across contexts (Figure S3B). Surprisingly, the hippocampal population exhibited this denser coactivity structure in context *Y* without a reduction in the average geodesic path length (see methods), calculated as the mean shortest path between any two nodes (Figure 2F) [8]. This suggested that the hippocampal population structure acquired in context *Y* exhibited the higher node-to-node coherence of a more regular, lattice-like network [9–11]. These topological alterations developed across conditioning days (Figure S3C) to continue altering the level and structure of population coactivity in post-conditioning days (Figure S3D), affecting the baseline re-exposure to context *Y* prior to any testing.

**Figure 2.**
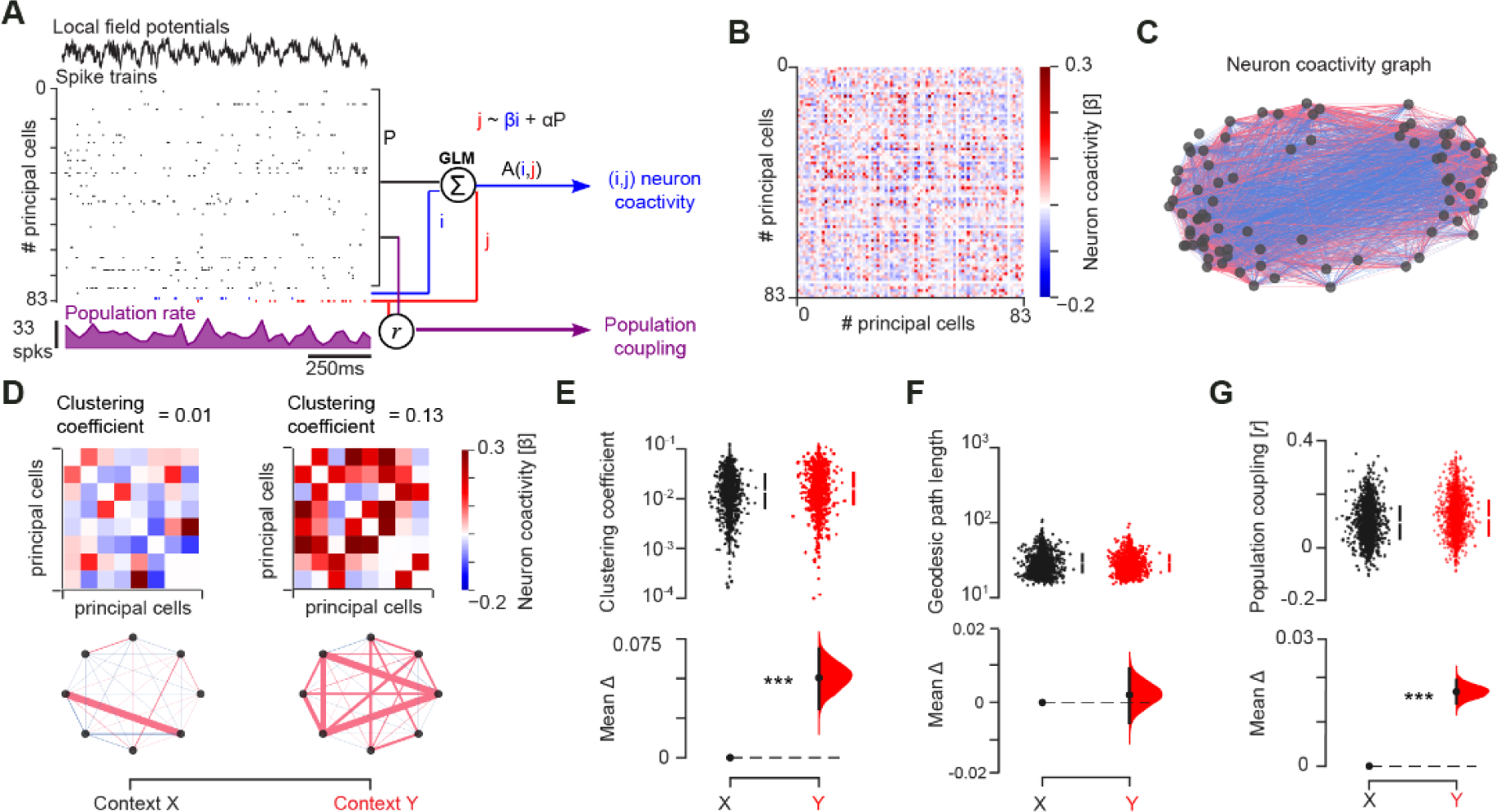
Robust memory increases neuronal coactivity and population coupling. **(A)** Schematic of population-level analyses (see methods). Coactivity between any two (*i*, *j*) neurons measured as the β regression weight from the GLM assessing their firing relationship while accounting for network-level modulation using the sum of the remaining cells in the population (to estimate neuron pair (*i*, *j*) coactivity beyond the population rate). Population coupling of each cell measured as the Pearson correlation coefficient between its theta-binned spike train and the cumulative activity of the remaining cells. **(B-D)** Example adjacency matrix of β regression weights (B) and corresponding coactivity graph (C) using the procedure depicted in (A) to access the neuron (*i*, *j*) pairwise coactivity structure of the population in each context. For clarity, (D) shows a subset of adjacency matrices representing contexts *X* versus *Y* (top), along with their clustering coefficients and motifs of coactivity (bottom). (**E, F)** Estimation plots showing that the population coactivity structure is tighter in context *Y* than *X*, as reported by the higher clustering coefficient of neuronal graphs containing more triads of coactive neurons (E), without a significant change in geodesic path length (F). **(G)** Population coupling of principal cells is stronger in context *Y* than *X*. ***P<0.001.

To further explore the development of a dense population activity structure in context *Y*, we investigated the one-to-many relationship between individual neurons and the rest of the population. For this, we measured across theta cycles the coupling of each principal cell firing to the summed activity of all other recorded cell members of the population (‘population rate;’ Figure 2A). Consistent with the higher topological clustering (Figure 2E), the average population coupling of individual neurons was stronger in context *Y* (Figure 2G). This increased population coupling reflected a stronger cross-neuron spiking relationship: shuffling the spike times across neurons and theta cycles, while preserving each neuron’s mean rate and the population rate distribution, cancelled the increased population coupling seen in context *Y* (Figure S4A). This heightened coupling developed across conditioning days to mark the re-exposure to context *Y* during post-conditioning days even before any test (Figure S4B). Contextual food conditioning thus seemed to increase the recruitment of principal cells as “choristers of a larger hippocampal orchestra’’ [12].

We therefore investigated the cellular substrate contributing to the denser correlational structure marking the hippocampal population after contextual food conditioning. We started by quantifying CA1 neuron recruitment in mice exposed to Hfd in context *Y* for 10 days, as indexed by the expression of the activity-dependent immediate-early-gene cFos. In parallel, control mice either ate Hfd in their homecage or explored context *Y* without food. Mice undergoing Hfd-context *Y* conditioning showed higher density of cFos^+^ neurons in the CA1 *pyramidale* layer compared to controls (Figure 3A, B). We further noted that 75.45 ± 3.52 % of these cFos-expressing cells were in the superficial *pyramidale* sublayer (i.e., adjacent to the *radiatum* layer), as revealed by the expression of the marker Calbindin 1 (Figure 3A) [13–15]. This suggested that contextual food conditioning recruited CA1 neurons preferentially in the superficial *pyramidale* sublayer.

**Figure 3.**
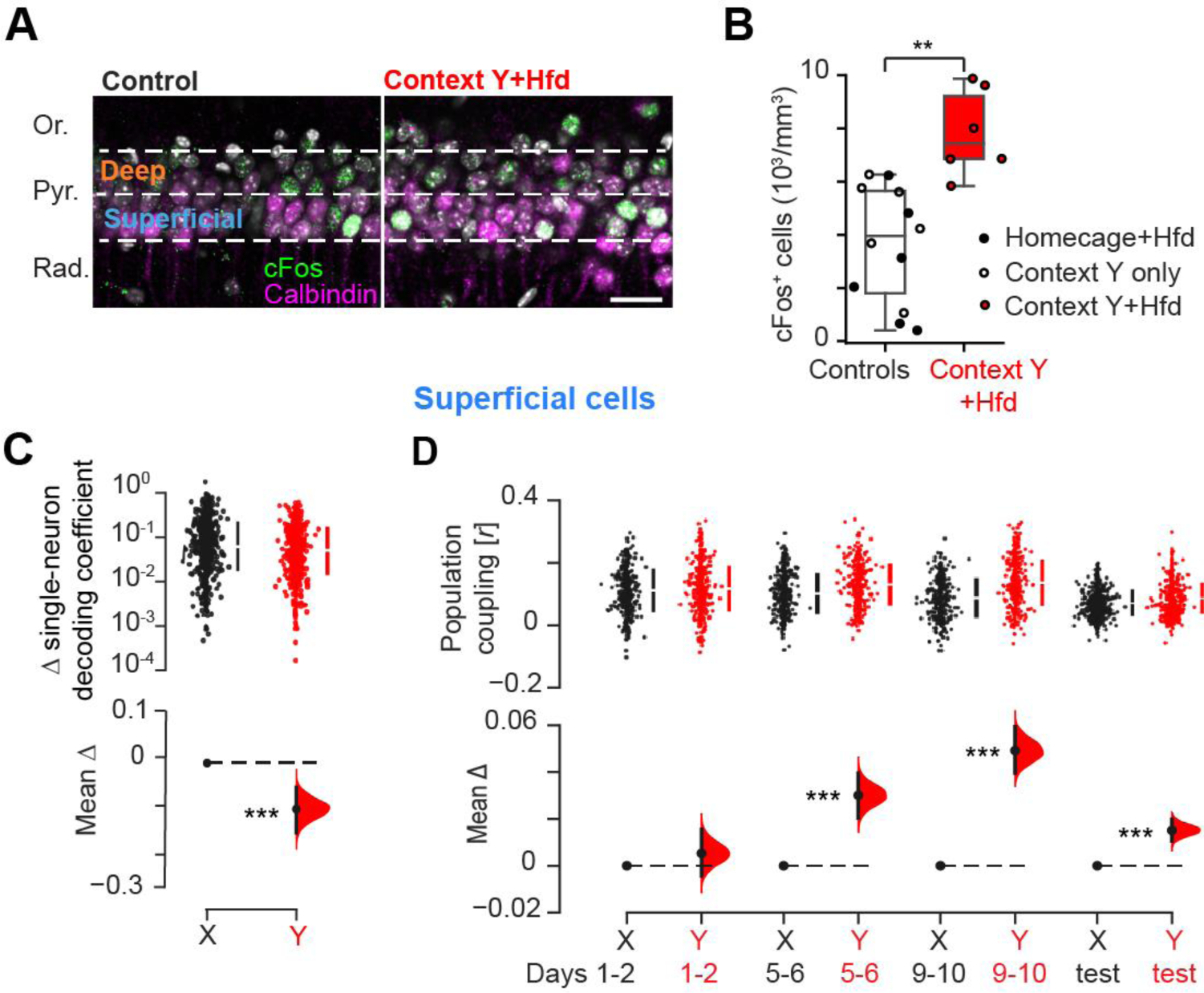
Contextual food memory recruits CA1 superficial *pyramidale* sublayer cells. **(A)** cFos–expressing neurons in the CA1 superficial *pyramidale* (Pyr.; *Oriens*, Or.; *Radiatum,* Rad.) sublayer (Calbindin 1 marker delineated) for a mouse after 10-day context *Y*-Hfd conditioning (right) and a control mouse not submitted to Hfd conditioning (left). Cell nuclei stained with DAPI (gray). Scale bar: 50 μm. (**B**) Corresponding quantification of cFos^+^ cell density in the CA1 pyramidal layer (each data point represents one mouse per condition; p=0.001, Kruskal-Wallis test). (**C, D**) Estimation plots showing the reduced change (update) in the GLM decoding contribution of single CA1 superficial *pyramidale* sublayer cells to population object-location classification (C; as in Figure 1J), and their increased coupling to the rest of the population (D) in context *Y* compared to *X*. ***P<0.001, **P<0.01.

We leveraged this observation by identifying neurons recorded in the CA1 superficial *pyramidale* sublayer to evaluate their contribution to memory. This identification was achieved using the electrophysiological profile of each tetrode (Figure S5) [8]. We found that the population representational rigidity that we had observed affecting object-location memory update across cNOR tests in context *Y* (Figure 1I, J) related to unchanged contribution of superficial cells. That is, superficial cells showed significantly reduced modulation of their individual contribution to the population GLM object decoding from cNOR session *n* − 1 to test *n* in context *Y* (Figure 3C). This result suggested that superficial cells did not seemingly distinguish the novel object from the familiar previously encountered at the same location in context *Y*, only effectively representing the location itself. Moreover, superficial cells developed stronger population coupling throughout conditioning in context *Y*, which persisted thereafter in test days (Figure 3D and Figure S6A). In line with this, superficial cells formed more triads of coactive neurons (Figure S6B, C). Cells located in the deep *pyramidale* sublayer did not show a similar activity profile (Figure S7). The CA1 representational rigidity preventing flexible processing of object-location information across memory tests in context *Y* was thus mostly explained by aberrant activity of superficial *pyramidale* sublayer cells.

We hypothesized that preventing the rise in population coupling during robust contextual conditioning would restore flexible memory. For this, we used an intersectional optogenetic strategy to target and repress cells recruited in the CA1 superficial *pyramidale* sublayer during Hfd-context *Y* conditioning. CA1 superficial *pyramidale* sublayer cells are genetically defined by the molecular marker Calbindin 1 [13,15,16]. They express cFos during contextual Hfd conditioning (Figure 3A, B). We thus bred double-transgenic Calb1-Cre;cFos-tTA mice and generated a viral construct for the two-term Boolean logic [17] expression of the yellow light-driven neural silencer ArchT-EYFP (or its EYFP-only control) dependent on the two recombinases Cre and FlpO (Figure 4A). We transduced the CA1 of these mice with this construct together with a second construct allowing the tTA-dependent expression of FlpO (Figure 4A) for lasting optogenetic tagging of CA1 superficial *pyramidale* sublayer cells with either ArchT-EYFP (in CA1^Calb1-cFos^::ArchT mice) or EYFP-only (in CA1^Calb1-cFos^::EYFP mice) from the onset of Hfd conditioning in context *Y* (Figure 4B and Figure S8A, B). CA1 superficial principal cells preferentially fire action potentials at the trough of theta cycles (Figure S8C) [8,18,19], further exhibiting increased theta modulation across conditioning days (Figure S8D). We thus combined our intersectional strategy with a closed-loop controller for real-time light-actuated suppression of CA1 superficial cells at their preferred theta phase during Hfd-context *Y* conditioning (Figure 4C-E and Figure S8A). We found that despite undergoing 10-day Hfd feeding in context *Y* (Figure S8E), this cell-type-defined network-pattern-informed intervention subsequently restored both natural food neophobia (Figure S8F) and successful novel object-location memory (Figure 4F) in the “optogenetically adjusted” CA1^Calb1-cFos^::ArchT mice. This was not the case for the “non-optogenetically adjusted” CA1^Calb1-cFos^::EYFP mice that overcame rodent food neophobia (Figure S8F) and showed impaired object-location memory (Figure 4F) in context *Y*, as our original mice (Figure 1D, E). In line with this behavioral outcome, optogenetically adjusted CA1^Calb1-cFos^::ArchT mice recovered functional object-location population decoding (Figure 4G) along with restored single-neuron representational flexibility (Figure S8G), population coactivity topology (Figure 4H, I) and coupling (Figure 4J) in both context *X* and *Y*. These findings showed that adjusting the hippocampal population coactivity structure in this manner supports adaptive transition from robust to flexible memory.

**Figure 4.**
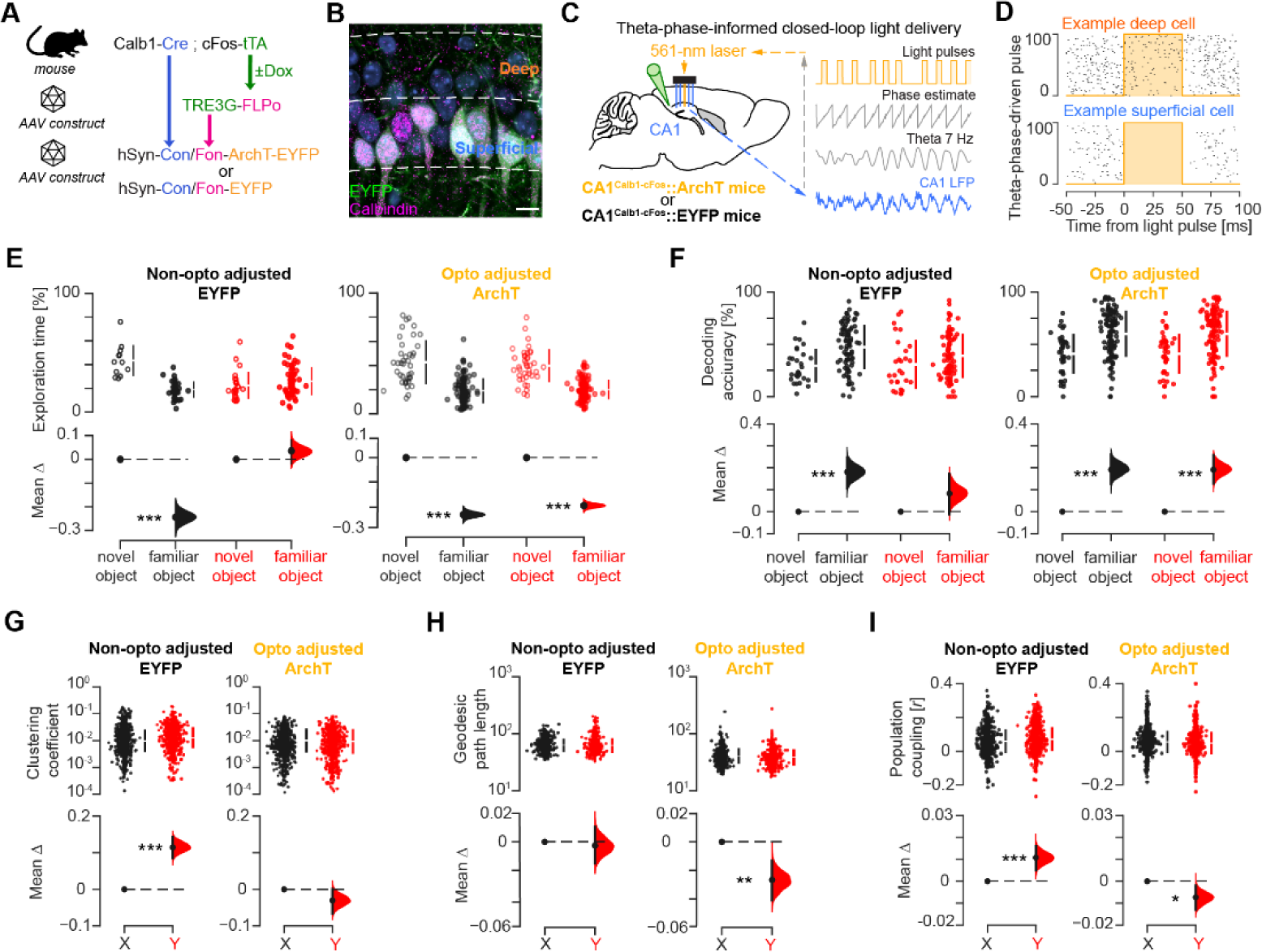
Adjusting hippocampal population coactivity restores flexible memory. **(A, B)** Intersectional optogenetic strategy (A) to target CA1 Calb1 neurons recruited during Hfd-context *Y* conditioning (B; *green*, EYFP; *magenta*, Calbindin 1; *blue*, DAPI). **(C, D)** Closed-loop CA1 light delivery controller (C) combined with the optogenetic strategy shown in (A) to suppress superficial *pyramidale* sublayer cells at their preferred theta phase (Figure S8C) during Hfd-context *Y* conditioning. (D) shows example spiking activity for a deep (top) versus a superficial (bottom) *pyramidale* sublayer cell with respect to theta phase-driven light delivery. **(E-I)** Estimation plots (see methods). For the optogenetically adjusted ArchT mice, but not the non-optogenetically adjusted EYFP-only mice, this cell-type-selective network-pattern-informed intervention restored normal cNOR performance (E) along with population object-location decoding (F), rebalanced coactivity topology (G, H) and population coupling (I) in context *Y*. ***P<0.001, **P<0.01, *P<0.05.

In conclusion, we found that a robust (food-context) memory raises the population coupling of CA1 superficial *pyramidale* sublayer neurons, creating a dense network coactivity structure. With respect to subsequent behavior, this acquired hippocampal topology of heightened coactivity relates to strong contextual (feeding) response and impaired novel information processing. Neural networks have been proposed to fall into the broad class of ‘small-world’ networks, a middle ground between regular and random networks where the combination of high clustering of elements (a property of regular networks) with short path lengths between elements (a property of random networks) would allow important properties of complex networks such as increased computational power and effective synchronizability [9,10,20–22]. Here, our observation that repeated food-context conditioning affects the coactivity structure of the network by increasing neuronal clustering without shortening node-to-node paths suggests that the hippocampal topology is deviating from a small-world toward a more coherent, regular lattice. This way, the joint activity of an increased number of neurons operating as a cohesive population would permit robust information flow to downstream receiver neurons, possibly at the expense of other (e.g., novel) input channels. This heightened hippocampal coactivity could be broadcasted to several recipient circuits and reader neurons. For instance, recent work points to a contribution for the nucleus accumbens in translating hippocampal dynamics of appetitive memory into a behavioral readout [23] or the hypothalamus in driving non-homeostatic contextual feeding [24]. Importantly, the instantiation of this highly clustered topology can be prevented: applying cell-type-selective, network-pattern-informed neuronal suppression during contextual learning rebalances hippocampal coactivity and restores flexible memory. Together, these findings suggest that a plastic organization of the hippocampal network topology constraining the collective activity of its neuron constituents shapes the integration of new memories into pre-existing knowledge, from behavioral adaptation to maladaptation and back.

## Acknowledgements

We would like to thank R. Lambiotte and S.R. Schultz for commenting on a previous version of the manuscript; B. Micklem for technical assistance; all members of the Dupret lab for feedback during the project. **Funding:** This work was supported by the Biotechnology and Biological Sciences Research Council UK (awards BB/S007741/1 and BB/N002547/1) and the Medical Research Council UK (programme MC_UU_00003/4 and award MR/W004860/1). **Author contributions:** Conceptualization, GPG, LL, and DD; Investigation, GPG, LL, TB, SBMc, DB, KH, HH; Analysis, GPG, LL, and DD; Methodology, GPG, LL, SBMc, VLdS, and DD; Resources, PVP, RT, AS, and DD; Visualization, GPG and DD; Funding acquisition: DD; Writing – Original Draft, GPG and DD; Writing – Reviewing & Editing, GPG, LL, TB, SBMc, VLdS, DB, KH, HH, PVP, RT, AS and DD; Supervision, DD. **Competing interests:** The authors declare no competing interests.

## Materials and Methods

### Animals

These experiments used adult (4–6 months old) C57BL/6J wild-type mice (Charles River Laboratories, UK) and double-transgenic mice obtained by crossing transgenic mice heterozygous for the transgene expressing the Cre recombinase under the control of the Calbindin 1 (Calb1) promoter (Jackson Laboratories; obtained from C57BL/6J crossed with Calb1-IRES2-Cre-D B6.129S-Calb1^tm2.1(cre)Hze^/J, stock number 028532, RRID: IMSR_JAX:028532) with c-fos–tTA transgenic male mice heterozygous for the transgene carrying the c-fos promoter–driven tetracycline transactivator (tTA)[25,26] (c-fos-tTA mouse line generated at The Scripps Research Institute, maintained at Tufts University until shipment to the MRC BNDU at the University of Oxford where they were bred from c-fos-tTA mice crossed with C57Bl6/J mice). Animals were housed with their littermates up until the start of the experiment. All mice held in IVCs, with wooden chew stick and nestlets in a dedicated housing facility with a 12/12 h light/dark cycle (lights on at 07:00), 19–23°C ambient temperature and 40–70% humidity. They had free access to water and food *ad libitum* throughout the experiment. Experimental procedures performed on mice in accordance with the Animals (Scientific Procedures) Act, 1986 (United Kingdom), with final ethical review by the Animals in Science Regulation Unit of the UK Home Office.

### Surgical procedure

All surgical procedures were performed under deep anesthesia using isoflurane (0.5–2%) and oxygen (2 l/min), with analgesia provided before (0.1 mg/kg vetergesic) and after (5 mg/kg metacam) surgery.

To generate expression of either ArchT-EYFP or EYFP-only in principal cells of the CA1 superficial *pyramidale* sublayer, we combined Cre-dependent, FlpO-dependent, and tTA-dependent approaches (see section “Viral constructs” below). We mixed in a 1:5 ratio a tTA-inducible AAV carrying a TRE3G-FlpO construct with a Cre-dependent FlpO-dependent AAV carrying either a hSyn-Con/Fon-ArchT-EYFP or the corresponding control hSyn-Con/Fon-EYFP. Viral injections were targeted bilaterally to dorsal CA1 hippocampus of double-transgenic Calb1-Cre;cFos-tTA mice (Figure 4 and Figure S8) using stereotaxic coordinates (-1.7 and -2.3 mm anteroposterior from bregma, ±1.25 and ±1.7 mm lateral from bregma, and -1.1 mm ventral from the brain surface; 150 nl per site; 2 injection sites per side) at a rate of 100 nl/min using a glass micropipette lowered to the target site and held in place for 5 min after virus delivery before being withdrawn.

For electrophysiological tetrode recordings, mice were implanted with a single microdrive containing 14 independently movable tetrodes, targeting the *stratum pyramidale* of the dorsal CA1 hippocampus [8]. Tetrodes were constructed by twisting together four insulated tungsten wires (12 μm diameter, California Fine Wire) which were briefly heated to bind them together into a single bundle. Each tetrode was loaded in one cannula attached to a 6 mm long M1.0 screw to enable its independent manipulation of depth. The drive was implanted under stereotaxic control in reference to bregma using central coordinates -2.0 mm anteroposterior from bregma, +1.7 mm lateral from bregma as a reference to position each individual tetrode contained in the microdrive, initially implanting tetrodes above the pyramidal layer (-1.0 mm ventral from brain surface). The distance between neighboring tetrodes was 350 μm. Following the implantation, the exposed parts of the tetrodes were covered with paraffin wax, after which the drive was secured to the skull using dental cement and stainless-steel anchor screws inserted into the skull. Two of the anchor screws, both above the cerebellum, were attached to a 50 µm tungsten wire (California Fine Wire) and served as ground. For the recordings, each tetrode was lowered along the vertical axis to reach the CA1 pyramidale layer, using the rotations applied to its tetrode cannula-holding screw and the electrophysiological profile of the local field potentials in the hippocampal ripple frequency band, with final depth position subsequently confirmed by histology of anatomical tracks.

For optogenetic manipulations, two optic fibers (230 μm diameter, Doric Lenses, Canada) were incorporated into the 14-tetrode microdrive designed to bilaterally deliver light to CA1 and implanted 10 days after viral injections.

In one mouse, a single-shank silicon probe (Neuronexus, model A1×32-5mm-25-177-H32_21mm) was implanted following the same surgical procedure to span the somato-dendritic axis of CA1 principal cells and establish the laminar profile of the sharp-wave ripples detected in the local field potentials. These silicon probe recordings allowed estimating the position (depth) of individual tetrode-recorded principal cell soma with respect to the deep versus the superficial sublayers of the dorsal CA1 *stratum pyramidale* (Figure S5).

For the hippocampal lesion surgery (Figure S2C, D), mice in the lesion group underwent the same anaesthetic induction as above, then scalp incision and craniotomy, followed by N-methy-D-aspartate (NMDA; 10 mg / mL) injections directly into the hippocampus at four sites per hemisphere using a modified Hamilton 36G syringe needle (anterior-posterior: -1.7, -2.3, -2.8, -3.1; mediolateral: ±1.2, ±1.7, ±2.2, ± 2.8; dorsoventral: -1.9, -1.9, -2.0, -4.0, respectively, 100 to 200 nL per site at the infusion / diffusion rates described above), and were then sutured. Midazolam (5 mg / kg, sub-cutaneous) was used to prevent seizures in hippocampal-lesioned mice. Mice receiving sham surgery were incised and then sutured. All mice had at least 2 weeks recovery before behavioural testing.

### Viral constructs

To optogenetically target cFos-expressing CA1 superficial *pyramidale* sublayer neurons genetically defined to express Calbindin 1, we first produced a TRE3G-FlpO AAV carrying the optimised FlpO recombinase under the control of the third generation of tetracycline responsive element containing promoter (TRE3G, Clontech Laboratories). For this, we exchanged ArchT-GFP open reading frame (ORF) (cut with NcoI and EcoRV) in pAAV-Tre3G-ArchT-GFP vector [26] with the Myc-FLPo ORF (cut with NcoI and Klenow blunted AscI) from pAAV-EF1a-DIO-FLPo-Myc vector [23] (Addgene plasmid # 124641; http://n2t.net/addgene:124641; RRID:Addgene_124641).

We next produced a viral vector allowing the Cre-dependent and Flp-dependent expression of ArchT-eYFP. The corresponding pAAV-hSyn-Con/Fon-ArchT-eYFP construct was cloned in two stages. First, pAAV-hSyn-Coff/Fon-ArchT-eYFP has been cloned. Plasmid vectors pAAV-CamKII-ArchT-GFP (a gift from Edward Boyden, Addgene plasmid #37807; http://n2t.net/addgene:37807; RRID: Addgene_37807)[27], pAAV-hSyn-Coff/Fon-hChR2(H134R)-eYFP (a gift from Karl Deisseroth, Addgene plasmid #55648; http://n2t.net/addgene:55648; RRID: Addgene_55648) [17] and the combination of the PCR products was used to assemble two inserts that were then subcloned into the pAAV-hSyn Coff/Fon hChR2(H134R)-eYFP vector to substitute the corresponding hChR2-eYFP coding exons with the ArchT-eYFP ones. Primers and the template plasmid DNA for the first insert (exon 1): GTTTCTGCTAGCAACCCCGACACTTACCTTAGCCAGCAGGGCCAG, GTTTCTGAGCTCGCCACCATGGACCCCATC, plasmid#37807. The PCR product was then cloned into the plasmid #55648 using NheI and SacI recognition sites thus forming the intermediate vector. For the following subcloning of the exon 2 the three PCR products were generated with the primers and the corresponding template DNAs: GTTTCTACTAGTCCTCCTGTACTCACC, GTGAGCAAGGGCGAGGAG, plasmid #55648; CTCCTCGCCCTTGCTCACTGCTACTACCGGTCGGGG, GACTCTATTTCTCATGTGTTTAGGTGGACAGGGTGAGCATCG, plasmid #37807; CCTAAACACATGAGAAATAGAGTC, CGAAGTTATGGTACCTGTGCCCCCCC, plasmid #55648. These three products were then combined in the overlapping PCR and inserted into the intermediate vector using SpeI and KpnI cloning sites forming pAAV-hSyn-Coff/Fon-ArchT-eYFP. pAAV-hSyn-Coff/Fon-ArchT-YFP vector then was used to produce pAAV-hSyn-Con/Fon-ArchT-eYFP by inverting the sequence containing part of ArchT-eYFP exon and flanked with the SpeI and KpnI restriction enzymes. The corresponding insert was produced by the PCR with the primers GTTTCTACTAGTTGTGCCCCCCCTTTTTTTTAT and GTTTCTGGTACCCCTCCTGTACTCACCTTGCC using pAAV-hSyn-Coff/Fon-ArchT-eYFP vector as a template.

The control vector expressing eYFP in the Cre-dependent and Flp-dependent manner was a gift from Karl Deisseroth (Addgene plasmid # 55650; http://n2t.net/addgene:55650; RRID:Addgene_55650) [17].

We mixed in a 1:5 ratio the AAV carrying TRE3G-FlpO and one of the two Cre-dependent Flp-dependent AAVs in the CA1 of double-transgenic Calb1-Cre;cFos-tTA mice (Figure 4 and Figure S8).

### Contextual feeding task

Following full recovery from the surgery, each mouse was first handled in a dedicated handling cloth, connected to the recording system, and exposed to an open-field enclosure (neither context *X* not *Y*) to be familiarized with the recording procedure over a period of one week prior to the start of the experiment itself. During this period, tetrodes were gradually lowered to the CA1 *stratum pyramidale* using their estimated depth location, local field potentials, and neuronal spike waveforms. Mice were always fed *ad libitum* throughout the entire experiment. Our contextual feeding task involved three open-field arenas referred to as context *X*, *Y*, and *Z*. These enclosures (outer dimensions: 46 x 46 cm; 38 cm height) differed in their shape (Figure 1A and Figure S1C) and wall-attached cue cards. In the first stage (“Conditioning”) of this task, mice underwent food-context conditioning over 10 consecutive days (Figure 1B; days 1–10). On each conditioning day, mice explored the arenas in a random order, for 30 minutes each. One arena (context *X*) always contained two food containers in each of which we provided one regular diet pellet (Chow; Figure S1A), identical to those present in the mouse homecage. Another arena (context *Y*) contained two similar food containers that allowed a choice between one chow pellet versus one high-fat diet pellet (Figure S1A; 45% Hfd; Research Diets Inc.; catalog number #D12451). The third arena (context *Z*) was not paired with any food resources during conditioning. On each conditioning day, mice were allowed to rest for 30 minutes between arena explorations.

The second stage (“Novel food test”) of this task both assessed changes in baseline population-level activity caused by the 10-day conditioning and probed discriminative food-context association. For these, on each of the two following days (days 11 and 12; Figure 1B), we first re-exposed mice to either context *X* or *Y* (“Re-exposure”; 30-min; Figure 1B) before we measured mouse propensity to express context-biased feeding (Figure S1B). In each novel food test, the arena contained two containers that allowed a choice between two food items: one had a regular chow pellet, the other had a never-seen-before resource (e.g., apricot or blackcurrant fruit jams; chocolate or peanut butter food pastes). The kCal value and palatability of novel food resources were matched across test days and contexts. Novel food intake was measured as the novel food kCal intake in context *Y* minus that in context *X* over their sum. A positive score thus indicates increased novel food intake in context *Y* compared to *X*. We further tested mouse propensity to eat Hfd in a context other than *Y* (“Hfd test”; Figure S1C). For this, mice explored for 30 minutes context *Z* that contained two food containers: one with a chow pellet, the other with a Hfd pellet. Hfd food intake was measured as the Hfd intake in context *Z* minus that measured in context *Y* in the previous day, over their sum.

We adapted this 12-day contextual feeding paradigm in Calb1-Cre;cFos-tTA mice in order to optogenetically tag the CA1 principal cells that are genetically defined by Calbindin 1 and further expressed cFos when recruited during Hfd conditioning in context *Y*. For this, Calb1-Cre;cFos-tTA mice were fed for 10 days with doxycycline-containing pellets (Envigo Ltd., Catalog number #TD120240) prior to viral construct injections and microdrive implantation (Figure S8A). In this set of experiments, we replaced the Chow pellets with Dox pellets in both context *X* and *Y*, except during the off-Dox tagging days in context *Y*. Before these tagging days, we conducted the first 2 days of conditioning in context *X* (Figure S8A; dox versus dox pellets). For optogenetic tagging, the dox-containing pellet homecage feeding was ceased for the first two days of conditioning in context *Y*, after which they received high-doxycycline containing pellets (Envigo Ltd., Catalog number #TD120658) as homecage feeding for 24 hours before returning to normal doxycycline containing pellets for the remaining of the experiment. During the 2 tagging days, mice only explored context *Y* with chow versus Hfd pellets (Figure S8A).

### continuous Novel Object Recognition task

On each cNOR task day (Figure 1B; days 13 to 16), mice first re-explored context *X* and *Y* (“Re-exposure”; without any food; *X* versus *Y* in random order across cNOR days; 15-min exploration). This re-exposure session evaluated lasting changes in baseline population-level activity. In the second context (*X* or *Y*) explored that day, we next inserted four novel objects (“Sampling”; 15-min exploration) per day). Mice continued to explore this context for four more sessions (“Tests”; 10-min exploration each cNOR test) that day (Figure S2A). Each object (outer dimension: 50 x 50 mm width; 55 mm height; example objects shown in Figure S2B) was each positioned midway along a given wall. From the sampling session to the first cNOR test, and then for each other cNOR test, we replaced one of the four objects last explored with a novel object. This procedure thus allowed the mouse to explore three familiar (previously seen) and one completely novel object on each cNOR test. On each test, we measured the time spent exploring each object and we calculated the percentage time spent with the novel object versus the (mean) percentage time spent with the familiar objects. The interval between two cNOR test sessions was 5-min, during which the mouse was in a sleep/rest box transiently placed within the recording arena.

### Multichannel data acquisition, position tracking and light delivery

The extracellular signals from each tetrode channel were amplified, multiplexed, and digitized using a single integrated circuit located on the head of the animal (RHD2164, Intan Technologies; http://intantech.com/products_RHD2000.html; pass band 0.09 Hz to 7.60 kHz). The amplified and filtered electrophysiological signals were digitized at 20 kHz and saved to disk along with the synchronization signals (transistor-transistor logic digital pulses) reporting the animal’s position tracking and laser activations. The location of the animal was tracked using three differently colored LED clusters attached to the electrode casing and captured at 39 frames per second by an overhead color camera (https://github.com/kevin-allen/positrack/wiki). The LFPs were down-sampled to 1,250 Hz for all subsequent analyses. For optogenetic intervention, a 561-nm diode pumped solid-state laser (Crystal Laser, model CL561-100; distributer: Laser 2000, Ringstead, UK) was used to deliver light bilaterally to the CA1 via a 2-channel rotary joint (Doric Lenses Inc.). This was performed using a closed-loop system to deliver in real time 561-nm light pulses in CA1^Calb1-cFos^::ArchT mice and CA1^Calb1-cFos^::EYFP mice using dynamic tracking of ongoing theta phase (Figure 4C,D). The real-time phase estimate was obtained using the OscillTrack algorithm (https://colinmcn.github.io/OscillTrack/) [28] implemented in the field-programmable gate array of the Intan Technologies interface board. Phase detection was obtained by continuously operating on the data stream coming from an input channel containing the CA1 LFPs. This input channel used as the phase reference was high-pass filtered using a 1st order digital infinite impulse response filter with a corner frequency of 0.4 Hz to remove amplifier offset and electrode drift, then down-sampled 125-fold to a rate of 160 Hz for processing. The phase estimation was operated with a loop-gain of 0.0625 at a centre frequency of 7 Hz. Stimulation was triggered with a phase lead, such that the target phase aligned with the middle of the 50-ms light pulse.

### Spike detection and unit isolation

Spike sorting and unit isolation were performed with an automated clustering pipeline using Kilosort (https://github.com/cortex-lab/KiloSort) via the SpikeForest framework (https://github.com/flatironinstitute/spikeforest) [29,30]. To apply KiloSort to data acquired using tetrodes, the algorithm restricted templates to channels within a given tetrode bundle, while masking all other recording channels. All sessions recorded on a given day were concatenated and cluster cut together to monitor cells throughout the experiment day. The resulting clusters were verified by the operator using cross-channel spike waveforms, auto-correlation histograms, and cross-correlation histograms. Each unit used for analyses showed throughout the entire recording day stable spike waveforms, clear refractory period in their auto-correlation histogram, and absence of refractory period in its cross-correlation histograms with the other units. Principal cells and interneurons were identified by their auto-correlograms, firing rates, and spike waveforms. In total, this study includes n = 4,423 CA1 principal cells: n = 2,506 for the original dataset (Figures 1, 2, and 3; n = 7 mice); and n = 910 CA1^Calb1-cFos^::ArchT neurons (n = 5 mice) versus n = 1,007 CA1^Calb1-cFos^::EYFP neurons (n = 4 mice) for the optogenetic dataset (Figure 4).

### Theta oscillations, sharp-wave ripples detection, and tetrode depth estimation

To detect theta oscillations from the LFPs, we applied Ensemble Empirical Mode Decomposition and selected bouts of at least 5 cycles during active exploratory behavior (animal speed > 2 cm/s) [31]. To detect sharp-wave ripples, the LFP signal of each CA1 pyramidal layer channel (for tetrode recordings) or recording site (for linear silicon probe recordings) were subtracted by the mean across all channels/sites (common average reference). These re-referenced signals were then filtered for the ripple band (110 to 250 Hz; 4th order Butterworth filter) and their envelopes (instantaneous amplitudes) were computed by means of the Hilbert transform. The peaks (local maxima) of the ripple band envelope signals above a threshold (5 times the median of the envelope values of that channel) were regarded as candidate events. Further, the onset and offset of each event were determined as the time points at which the ripple envelope decayed below half of the detection threshold. Candidate events passing the following criteria were determined as SWR events: (*i*) ripple band power in the event channel was at least 2 times the ripple band power in the common average reference (to eliminate common high frequency noise); (*ii*) an event had at least four ripple cycles (to eliminate events that were too brief); (*iii*) ripple band power was at least 2 times higher than the supra-ripple band defined as 200-500 Hz (to eliminate high frequency noise, not spectrally compact at the ripple band, such as spike leakage artefacts). In addition to the SWR profile information, we integrated mean theta profile information to estimate the tetrode depth. We classified tetrodes as being in the deep or superficial sublayer of the CA1 *stratum pyramidale* based on the mean profile of detected SWRs and theta cycles (Figure S5). Positive values indicated that the tetrode was in the deep sublayer (i.e. closest to *stratum oriens*) while negative values indicated tetrode was in the superficial sublayer (i.e. closest to *stratum radiatum*) [8].

### Object-location population decoding

We trained a General Linear Model (GLM) to classify each object-location compound explored during cNOR tests, from the recorded neural spiking activity nested in theta cycles associated with these visits (Figure 1F). By training a model in session *n* − 1 and testing it in test *n* we then obtain one novel and three familiar object scores per cNOR session (four session pairs per day; Figure 1G). A given object-location compound pair *i* was then associated to a decoding vector composed of each neuron’s GLM coefficient (*P*^*i*^), reporting the individual weight contribution of neuron *n* to the population representation of *i* (Figure 1H). By taking the cosine distance metric between pairs of decoding vectors across subsequent sessions representing the same location but different objects (familiar object *i* versus novel object *j*), we quantified the change in population representation *D*_*pop*_ = *dist_cos_*(*P*^*i*^, *P*^*j*^) (decoding vectors similarity (Figure 1I). The representational change of individual neurons induced by the insertion of novel objects, was instead computed as 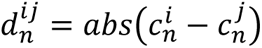 (Δ single-neuron decoding coefficient, Figure 1J). To build the GLM model, we employed the function *LogisticRegression* from the *sklearn* package (v .0.18…) with L2 regularization (inverse strength *C*=1e-2, selected by maximizing the classification scores across sessions).

### Neuronal coactivity graphs

We constructed hippocampal population graphs that represent the coactivity relationships between all pairs of principal cell spike trains recorded during a given task session. These coactivity graphs were computed using theta cycles as time windows spanning active exploratory behavior (speed>2 cm.s-1), discarding periods of immobility and further excluding sharp-wave/ripples. To further control for the shared influence of the general network activity on peer-to-peer coactivity, we used for any two neurons (*i*, *j*) the regression coefficient β _*ij*_ obtained by fitting the GLM (Figure 2A):

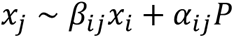

where *x*_*j*_, *x*_*i*_ are the z-scored theta-nested spike trains of individual neurons *j* (the target) and *i* (the predictor), and *P* is the summed activity of the other *N* − 2 neurons,

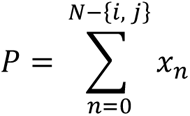

with *α*_*ij*_ weighting the influence of the population contribution to the activity of target neuron *j.* The recorded neurons (and their coactivity associations) are therefore the nodes (and their edges) in the coactivity graph of each task session. We described each graph by its adjacency matrix, *A*, as the *N* x *N* square matrix containing the pairwise coactivity relations within the network, yielding a weighted graph with no self-connections:

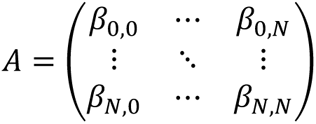

with β_*i*,*i*_ = 0 ∀*i in N*, and the symmetry in the weights of the network being ensured by setting 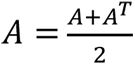 to form an undirected graph.

### Clustering coefficient

We computed a clustering coefficient to characterize the local synchronization of network activity by quantifying the number of three-node motifs. In each graph, for any neuron *i*, we obtained its clustering coefficient *C*_*i*_ using the formula proposed by Onnela et al. to quantify the strength of each triad [32–34]:

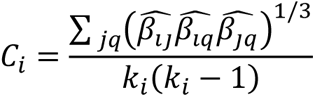

where *j* and *q* are neighbors of neuron *i*, all edge weights are normalized by the maximum edge weight in the network 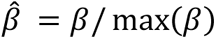, and *k_i_* is the degree of neuron *i*, which in these weighted graphs with no self-connection is equal to the number of neurons minus one. Note that this formula accounts for negative edges, yielding a negative value when there is an odd number due to the negative edges in the triad; it is positive otherwise. For this reason, this quantity can also be interpreted as a measure of the “structural balance” around a node, in the sense that its neighborhood presents coherent patterns of connectivity [35].

### Geodesic path length

We measured the geodesic (i.e., shortest) path length to estimate the activity coordination efficacy between any two nodes in the coactivity graph. In a binary graph, this would represent the smallest number of edges connecting two nodes. Here, we define the length between two nodes *i* and *j* as the inverse of their coactivity: *l* = 1/β_*ij*_, discarding all negative edges [11]. We then computed the geodesic shortest path length between any two nodes in the graph using the Floyd-Warshall algorithm [36–38]. We then report the average path length for each principal cell in the coactivity graph.

### Single-neuron coactivity strength

We defined the single-neuron coactivity strength as the average pairwise activity correlation strength of a given node in a weighted graph. As a reference, the strength in a weighted graph can be compared to the degree in a binary graph, which accounts for the number of the node’s neighbors. Here, the strength *S*_*i*_of a node *i* is the average across all the weights β_*ij*_ of the edges projected from that node:

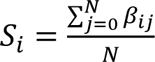

where *N* is the number of neurons *j* that node *i* projects to.

### Population coupling

We defined the coupling of neuron *i* to the rest of the population of *N* units (population coupling, *PC*) as the Pearson correlation between its theta-binned activity *x*_*i*_ and that of all remaining neurons summed together *P*^*N*−{*i*}^:

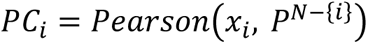

To assess whether the increase in population coupling in context *Y* was due to the fine temporal structure of the neural population activity, we independently circularly shuffled the neurons’ theta-binned activity by random delays between 10 to 100 theta cycles, therefore preserving each neurons’ rate and autocorrelation. The final value was taken as the average over 100 iterations.

### Anatomy

Mice were deeply anesthetized with isoflurane/pentobarbital and transcardially perfused with PBS followed by cold 4 % PFA dissolved in PBS. The brains were extracted, kept in 4% PFA for 24 h, and sliced into 50 µm thick coronal sections. For immunostaining, free-floating sections were rinsed extensively in PBS with 0.25 % Triton X-100 (PBS-T) and blocked for 1 h at room temperature in PBS-T with 10 % normal donkey serum (NDS). Sections were then incubated at 4°C for 48 h with primary antibodies (rabbit anti-cFos, 1:500, Synaptic Systems, cat# 226 003; goat anti-calbindin 1, 1:1,000, Nittobo Medical, cat# Af1040) diluted in PBS-T with 3 % NDS blocking solution. Sections were then rinsed three times for 10 min in PBS and incubated for 24 h at 4°C in secondary antibodies (donkey anti-rabbit Alexa Fluor 488, 1:250, Thermo Fisher Scientific, cat# A-21206, RRID:AB_2535792; donkey anti-goat Cy3, 1:1000, Jackson ImmunoResearch, cat# 705-165-147, RRID: AB_2307351) diluted in PBS-T with 1 % NDS. This step was followed by three rinses for 15 min in PBS. Sections were then incubated for 1 min with 4’,6-diamidino-2-phenylindole (DAPI; 0.5 μg/ml, Sigma-Aldrich, cat# D8417) diluted in PBS to label cell nuclei, before undergoing three additional rinse steps of 10 min each in PBS. Sections were finally mounted on slides, cover-slipped with Vectashield mounting medium (Vector Laboratories) and stored at 4 °C. Sections were also used for anatomical verification of the tetrode tracks. The CA1 pyramidal layer was imaged in all 3 channels (DAPI-A405, cFos-A488, and Calb1-CY3) as a mosaic stack across the full z-range using a Zeiss confocal microscope (LSM 880 Indimo, Axio Imager 2) with a Plan-Apochromat 20x/0.8 M27 Objective and the ZEN software (Zeiss Black 2.3, step size: 10μm) using 2 sections at the same rostro-caudal position (-2.0mm and -2.5 mm) for each mouse. Automated counting was performed using Fiji and the 3D ImageJ Suite [39]. A CA1 stratum pyramidale mask was generated using the DAPI channel, subdividing it into the deep versus the superficial pyramidale sublayers using the Calbindin 1 channel. To automatically detect single cFos cells, a 3D iterative thresholding (minimum threshold: RenyEntropy, pixel-volume: 2000-3000) was used on the cFos stack. A cFos positive cell was also positive for Calb1 when the average intensity within the cFos segment was above the automatic threshold (3D intensity measure, threshold: Huang). Cell densities are number of cells divided by the volume of the pyramidale layer. For the hippocampal lesion experiment, mice were perfused transcardially with physiological saline (0.9% NaCl) followed by 10% formol saline (10% formalin in physiological saline). The brains were then removed and placed in 10% formol saline and 72 hours later transferred to 30% sucrose-formalin. Coronal sections (50 μm) were cut on a freezing microtome and stained with cresyl violet to enable visualization of lesion extent.

### Data and statistical analyses

Data were analysed in Python 3.6 and using the packages DABEST v0.3.0 [40], scikit-learn v0.23.2 [41], NetworkX v2.4 [42], Numpy v1.18.1, Scipy v1.4.1, Matplotlib v3.1.2, Pandas v0.25.3 and Seaborn v0.11.0. All statistical tests related to a symmetric distribution were performed two-sided using Gardner-Altman plots (to compare 2 groups) and Cumming plots (for more groups) from the Data Analysis with Bootstrap-coupled ESTimation (DABEST) framework [40]. These DABEST plots allow visualizing the effect size by plotting the data as the mean or median difference between one of the groups (the left-most group of each plot, used as group-reference) and the other groups (to the right, along the x-axis of each plot). For each estimation plot: (*i*) the upper panel shows the distribution of raw data points for the entire dataset, superimposed on bar-plots reporting group mean±SEM, unless stated otherwise; and (*ii*) the lower panel displays the difference between a given group and the (left-most) group-reference, computed from 5,000 bootstrapped resamples and with difference-axis origin aligned to the mean or the median of the group-reference distribution. For each estimation plot: *black-dot*, mean (for normal distributions) or median (for skewed distributions) as indicated; *black-ticks*, 95% or 99% confidence interval as indicated; *filled-curve*: bootstrapped sampling-error distribution. Data distributions were assumed to be normal, but this was not formally tested. We also used the t-test to compare two conditions; the Wald test for assessing the significance of regression lines; and the Kolmogorov– Smirnov test for comparing probability distributions. Neural and behavioural data analyses were conducted in an identical way regardless of the identity of the experimental condition from which the data were collected, with the investigator blind to group allocation during data collection and/or analysis.

---

**Figure S1.**
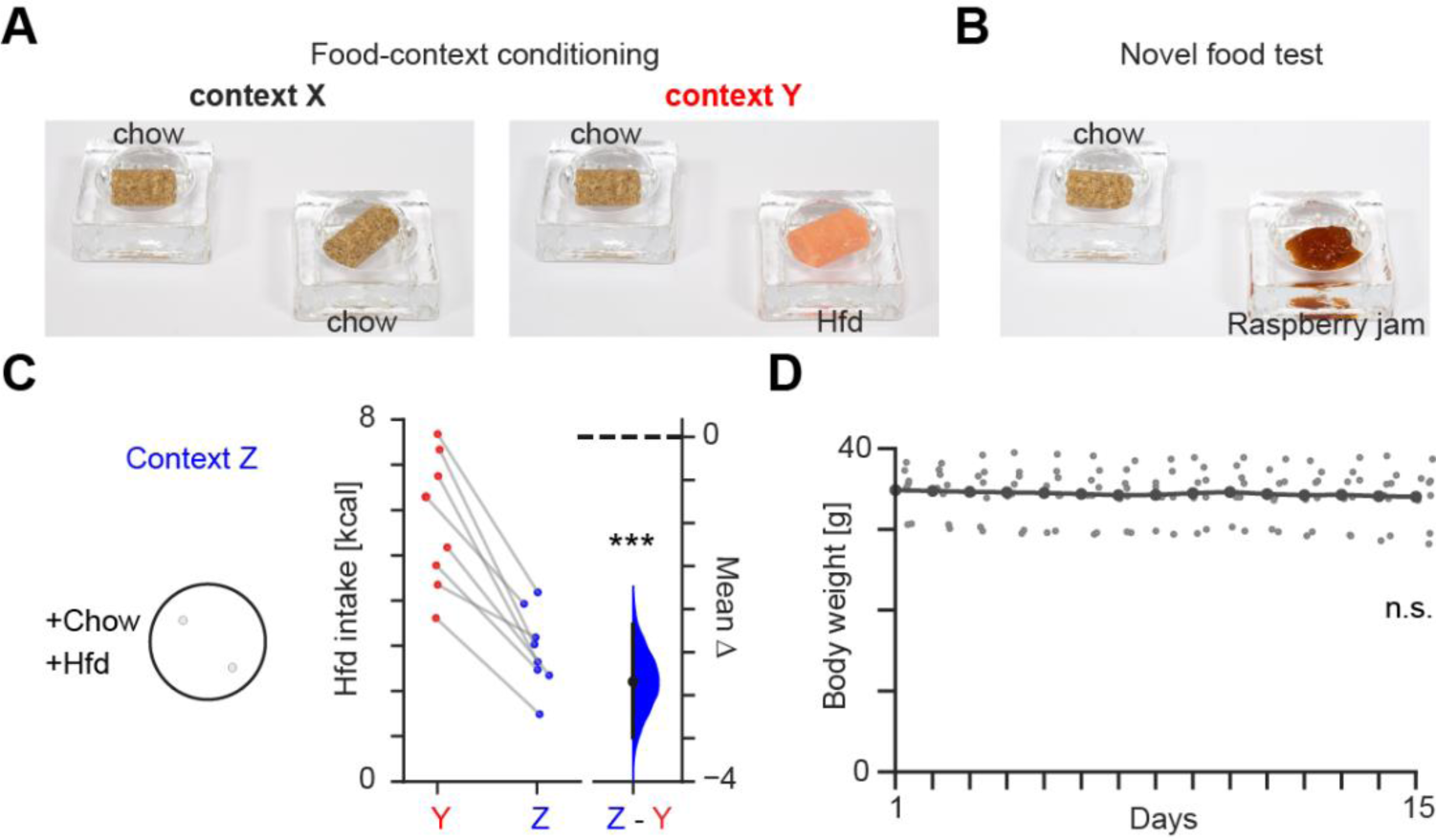
Contextual feeding task (days 1–12). **(A)** Pictures of the two food containers used to provide mice with a choice of two chow pellets in context *X* (left) and a choice between one chow pellet versus one high-fat diet (Hfd) pellet in context *Y* (right). These were used on each day of the food-context conditioning (days 1 – 10). **(B)** Likewise, shown are the two food containers used to provide mice with a choice between one chow pellet versus a new food resource (e.g., raspberry jam) during the post-conditioning novel food tests (days 11 and 12). **(C)** Layout and intake for the post-conditioning Hfd food test. Shown on the left is a top-view schematic of context *Z* as a circular-shaped arena that mice also explored from day 1 to day 10, without any food. For the post-conditioning Hfd food test, mice re-explored context *Z* with two food containers providing a choice between one chow pellet versus one Hfd pellet (as in context *Y* during conditioning; see A). Shown on the right is the corresponding estimation plot (see methods) showing the effect size for the difference in Hfd intake between context *Y* versus context *Z*. Note the significantly stronger Hfd food intake in context *Y* compared to *Z*. ***P<0.001. **(D)** Mice body weight across experiment days remained stable (each data point represents one mouse; t=-0.68, p=0.5; multiple regression).

**Figure S2.**
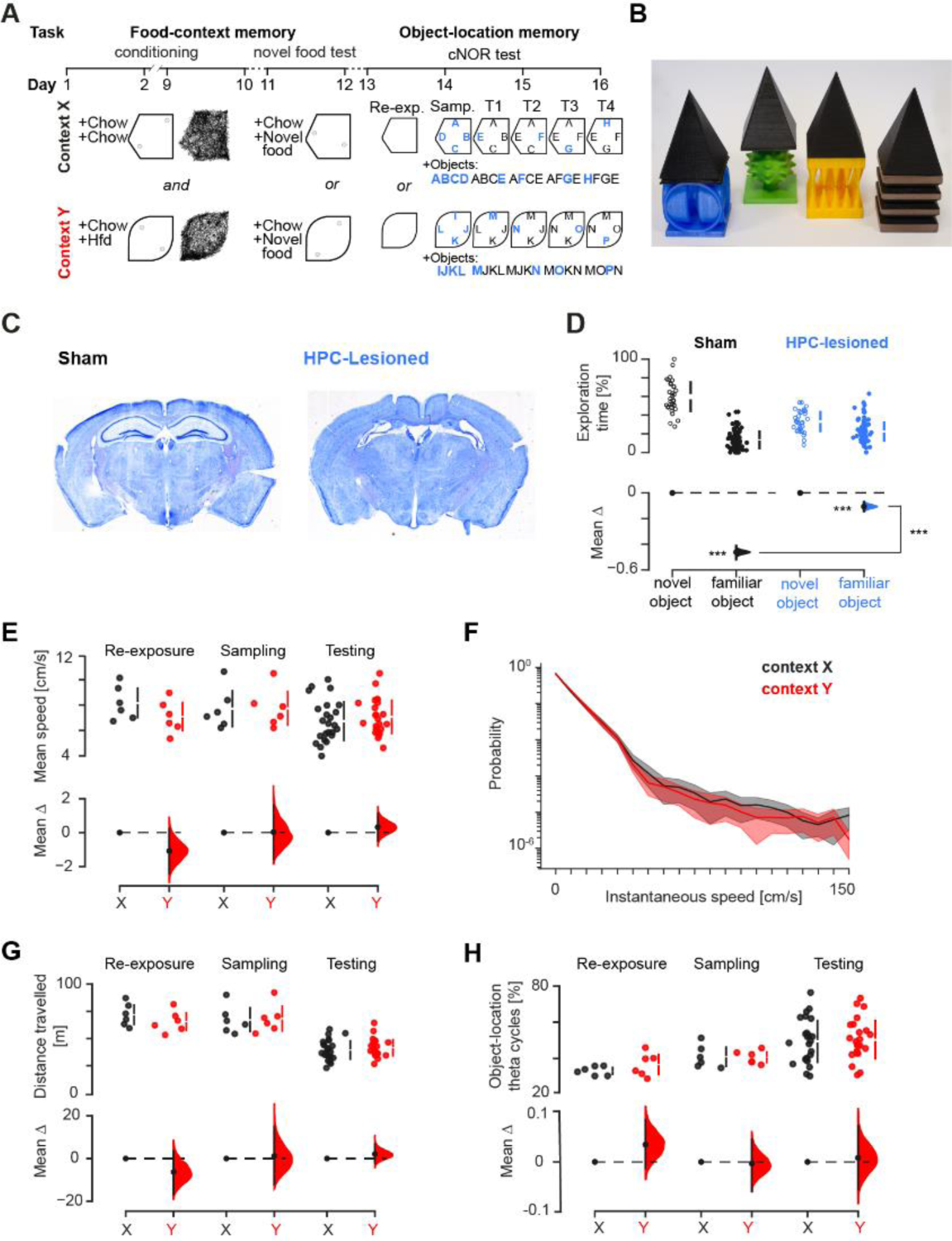
Continuous Novel Object Recognition task (days 13–16). **(A)** Layout of the 16-day experimental paradigm (as in Figure 1B) including further details about the continuous Novel Object Recognition (cNOR) task (days 13–16). On each cNOR day, we first re-exposed mice to context *X* and *Y* (Re-exposure labelled “Re-exp.”; 15-min exploration without any food; *X* and *Y* explored in random order across days) to evaluate post-conditioning changes in baseline population-level activity. Then, in the second context (*X* or *Y*) explored that day, mice explored four novel objects (Sampling labelled “Samp.”; 15-min exploration). Each object was positioned midway along a given wall. Over four additional exploration sessions (Tests labelled “T1 to T4”; 10-min each), mice continued to explore this context where one of the initial objects from the sampling session was replaced by a novel one. That is, by the fourth cNOR test, all initial novel objects (e.g., A, B, C, and D) were replaced by another one (e.g., E, F, G, and H). Using this procedure, mice were exposed to three familiar (previously seen) and one completely novel object on each cNOR test. On each test, we measured the time spent on each object and we calculated the percentage time spent investigating the novel object versus the (mean) percentage time spent investigating the familiar objects. **(B)** Example set of four objects. The “hat” on each object was used to prevent mice from climbing and staying onto the objects. (**C, D**) Hippocampal lesions impair cNOR performance. Shown in (C) is an example histology of coronal brain sections from one sham-lesioned and one hippocampal lesioned mouse (ibotenate infusion). Shown in (D) is an estimation plot for the effect size in the difference in time spent exploring novel versus familiar objects. Mice with sham lesions showed a strong preference for the novel object over the familiar (p < 0.001, two-sided paired permutation test). Mice with hippocampal lesions showed significantly weaker cNOR performance (p < 0.001, two-sided paired permutation test). Direct comparison showed significantly stronger novelty preference in sham compared to hippocampal-lesioned mice (p = 0.005, two-sided permutation test). (**E-G**) Locomotor speed and distance travelled did not differ between context *X* and *Y*. (E) is an estimation plot showing that the mean speed of the animals on each cNOR task days remained similar across the re-exposure, sampling, and test sessions in context *Y* compared to context *X*. This is also reported in the distribution of instantaneous speed values (F) and the total distance travelled (G). (**H**) Estimation plot showing that the percentage of theta cycles during exploration of each object-location was similar across contexts. ***P<0.001.

**Figure S3.**
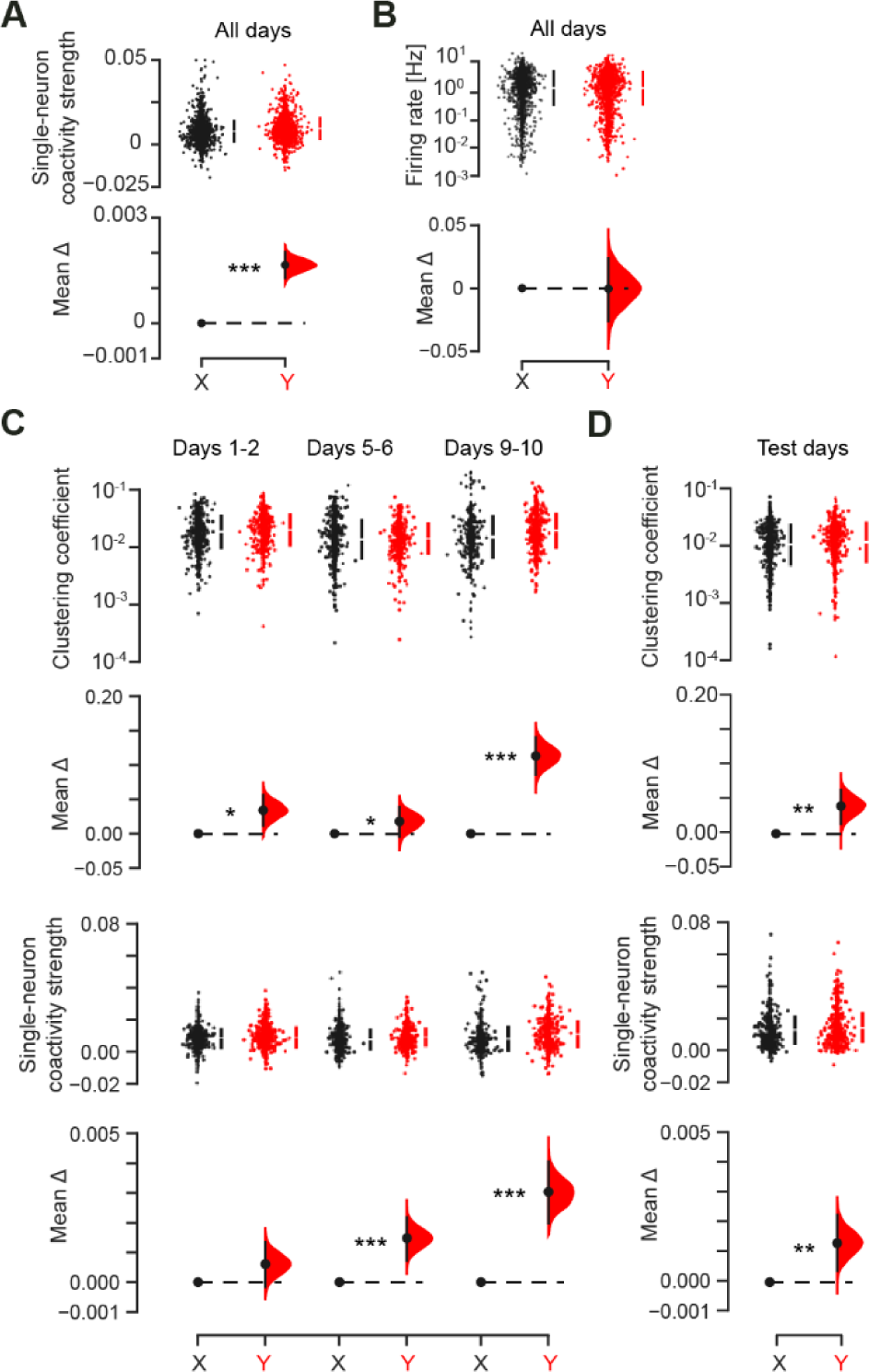
The topological changes affecting hippocampal neuronal graphs are developing during conditioning and are maintained thereafter. **(A, B)** Estimation plots showing stronger population coactivity level in context *Y* than *X*, as reported by higher mean pairwise coactivity strength (A), while the mean firing rate of CA1 principal cells was not different across contexts (B). All conditioning and test days together. See also Figure 2E for the mean clustering coefficient. **(C, D)** The mean clustering coefficient (top) and coactivity strength (bottom) increased in context *Y* compared to *X* over the conditioning days (C), to continue affecting neuronal graphs during the re-exposure session in post-conditioning test days (D). ***P<0.001, **P<0.01, *P<0.05.

**Figure S4.**
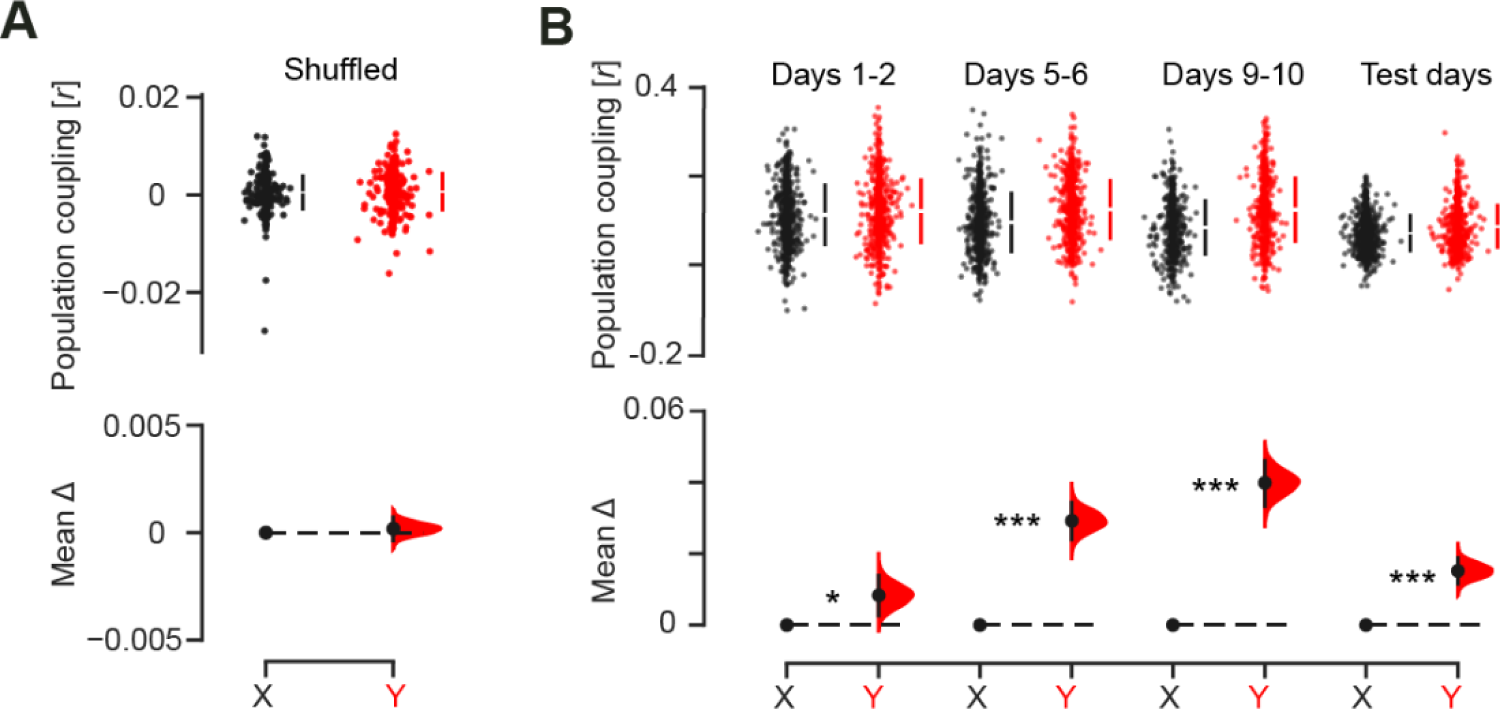
The population coupling of individual neurons is increasing during conditioning and is maintained thereafter. **(A)** Shuffling the spike times across neurons and theta cycles, while preserving each neuron’s mean rate and the population rate distribution, cancelled the increased average population coupling of individual neurons seen in context *Y* (Figure 2F). For this control analysis, the activity of each neuron was circularly shuffled independently with a delay drawn from a uniform distribution between 10 and 100 theta cycles. This result indicates that the increased population coupling seen in context *Y* compared to *X* (Figure 2F) reflected stronger cross-neuron spiking relationship. **(B)** This heightened population coupling developed across conditioning days to continue marking the re-exposure to context *Y* before any test during post-conditioning days. ***P<0.001, *P<0.05.

**Figure S5.**
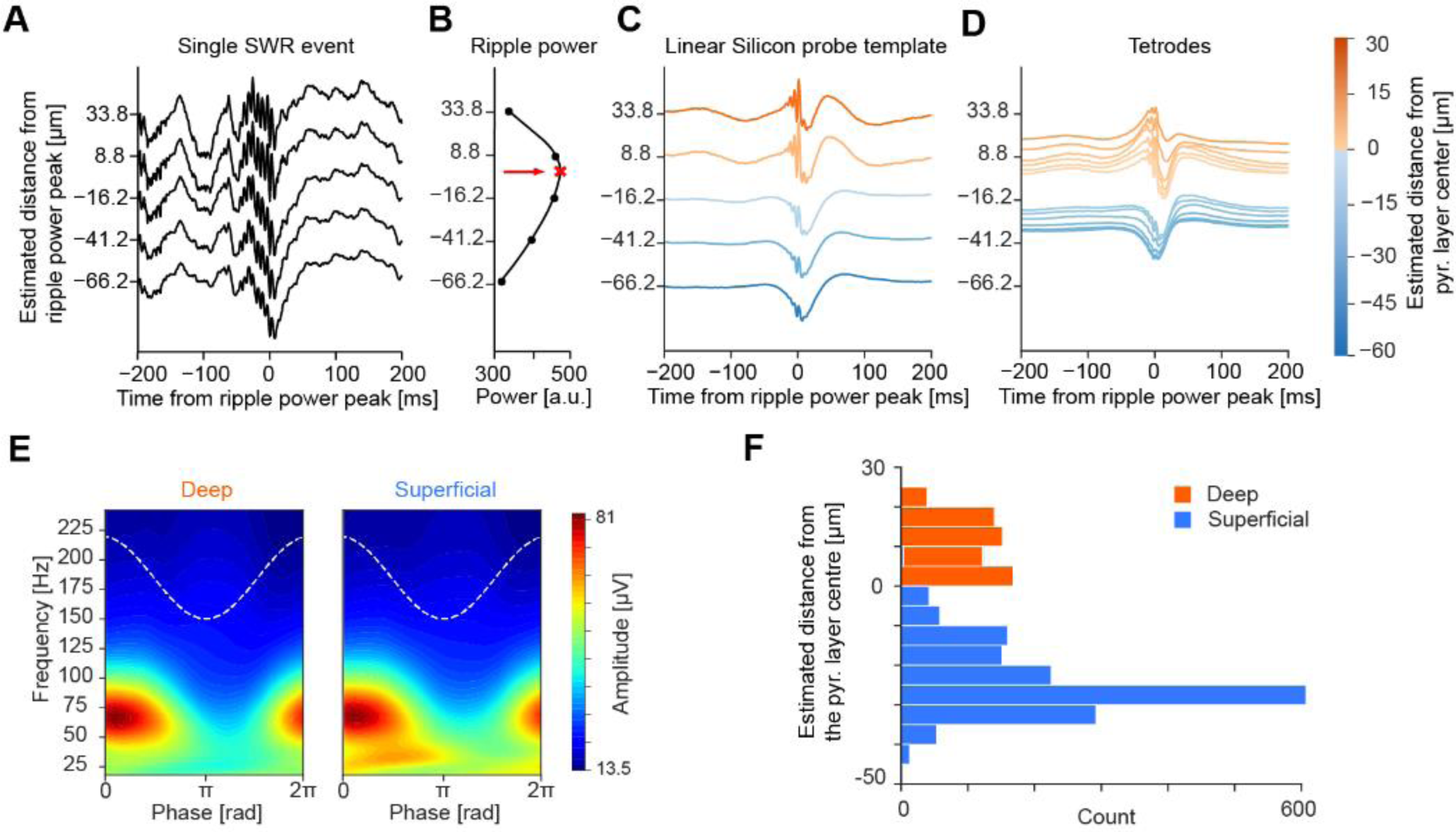
Identification of deep versus superficial CA1 *pyramidale* sublayer cells. **(A-D)** We estimated the position (depth) of individual tetrode-recorded principal cell soma by leveraging silicon probe recordings with known spacing between the recording sites along a linear shank (25-μm steps) [8]. From these silicon probe recordings, we first acquired the laminar profile of sharp-wave ripples (SWRs) detected in the LFPs along the radial axis of the CA1 hippocampus (A). We used the peak of the corresponding depth profile of the ripple-band (110–250Hz) power to estimate the centre (middle) of the pyramidal (pyr.) layer (B; red cross indicated with red arrow). Using the average LFP waveform of the SWR events detected in these silicon probe recordings, we then established a SWR template where we reported the distance relative to the estimated centre of the pyramidal layer, knowing the precise distance between the recording sites on the linear shank (C). Next, we computed the SWR profile of each individual tetrode to estimate its depth (and thus that its recorded neurons) by positioning its SWR profile within the silicon probe SWR template (the ground-truth vertical depth; C). Shown in (D) are examples of single-tetrode SWR profiles and their estimated depth. The SWR information was further integrated with the theta oscillation profile. **(E)** We further computed the average spectral profile for deep and superficial *pyramidale* tetrodes. This independent analysis showed that the spectral content of the LFP signals recorded from deep and superficial tetrodes differed with respect to the presence of slow gamma oscillations. **(F)** Corresponding distribution of principal cells (deep sublayer, n=635; superficial sublayer, n=1874) with respect to the estimated depth of their recording tetrodes.

**Figure S6.**
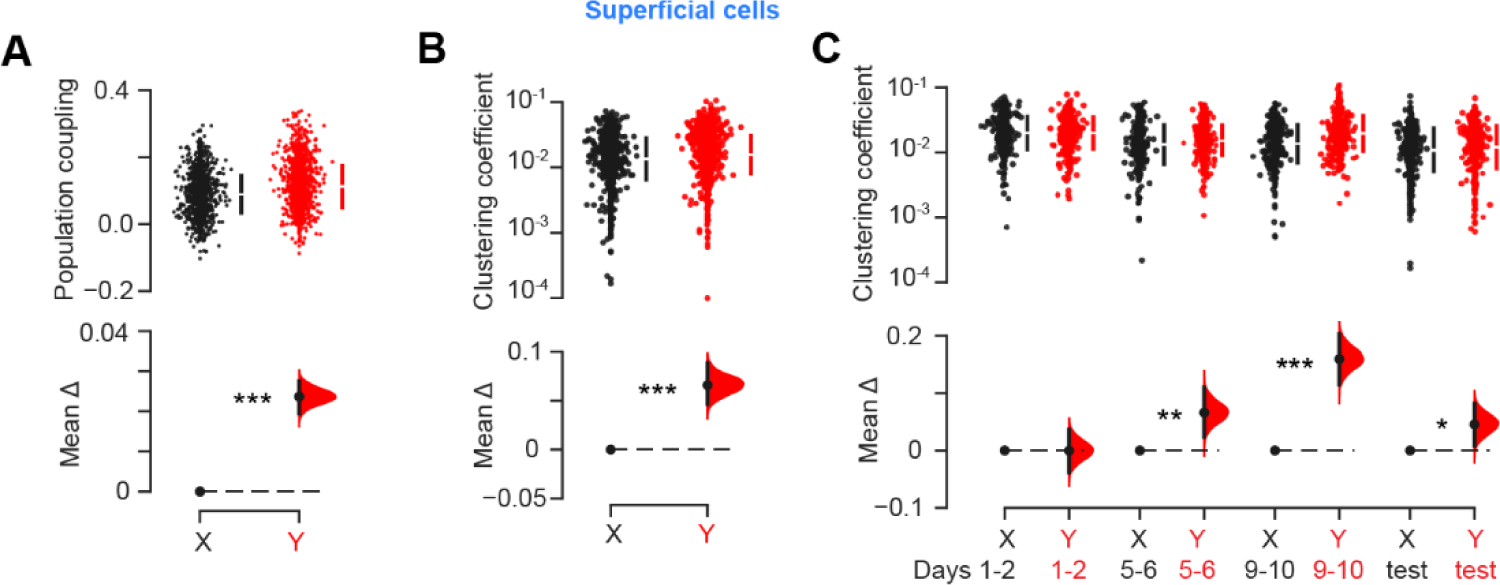
Principal cells in the superficial sublayer of CA1 *stratum pyramidale* increase their coactivity with contextual food conditioning. **(A, B)** Both population coupling (A) and coactivity clustering coefficient (B) of individual superficial CA1 *pyramidale* sublayer neurons are stronger in the Hfd-paired context *Y* compared to the chow-paired context *X*. All conditioning and test days together. (**C)** In line with this observation, the coactivity clustering coefficient of superficial cells increased in context *Y* across task days from conditioning to testing (as their population coupling; Figure 3E). ***P<0.001, **P<0.01, *P<0.05.

**Figure S7.**
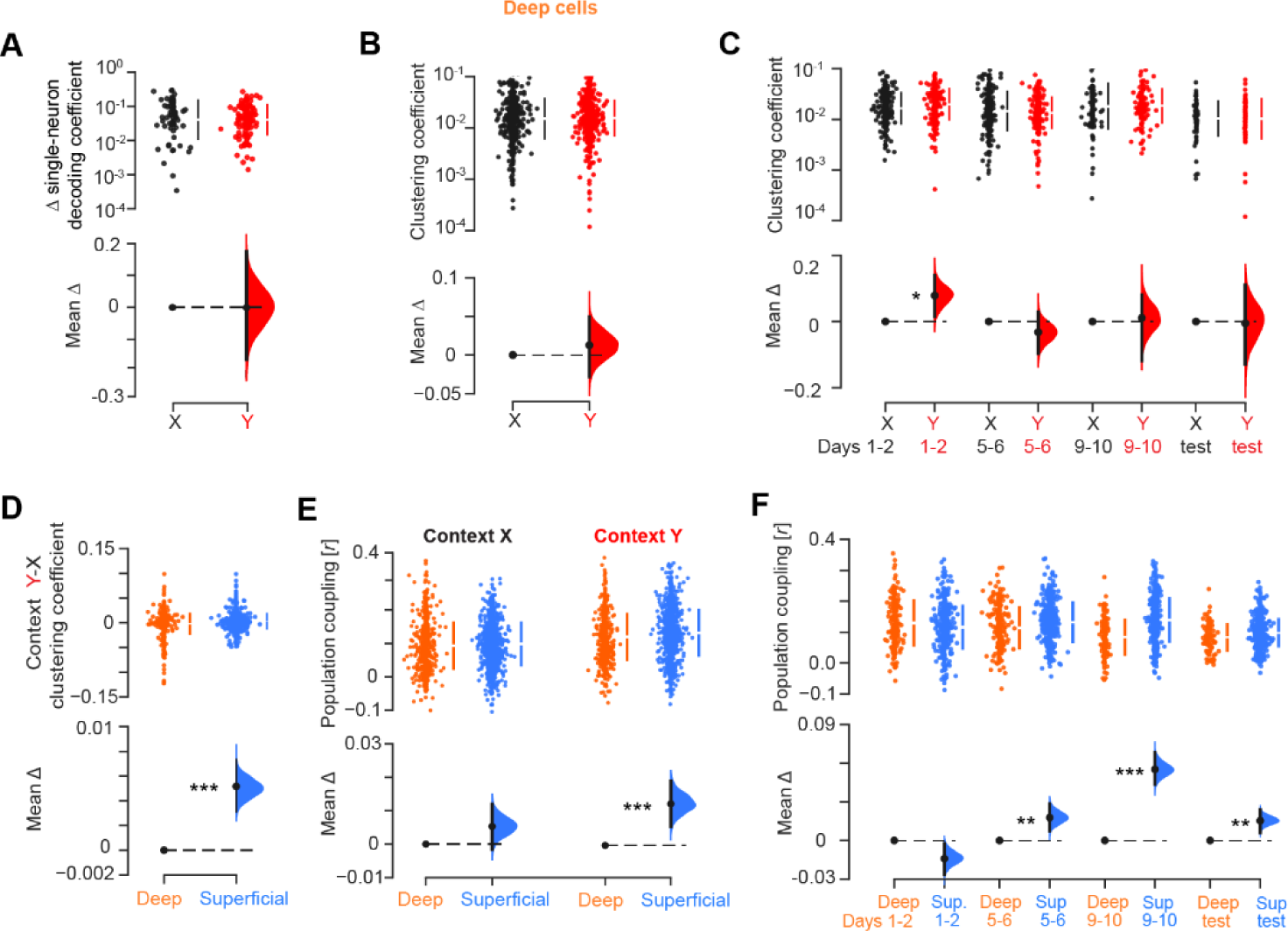
Deep versus superficial CA1 *pyramidale* sublayer cell activity. **(A)** Estimation plot showing the change in the magnitude of single-neuron classification contribution to novel object-location population decoding for CA1 deep *pyramidale* sublayer cells, quantified as the difference in their GLM coefficients in either context *X* or Y (see also Figure 3C and methods). **(B)** Estimation plot showing no change in coactivity clustering coefficient for deep *pyramidale* sublayer cells between context *X* and *Y*. All conditioning and test days together. **(C)** In line with this observation, the coactivity clustering coefficient of deep *pyramidale* sublayer cells remained constant in context *Y* compared to *X* from conditioning to testing. **(D)** Estimation plot showing the change in coactivity clustering coefficient for CA1 deep versus superficial *pyramidale* sublayer cells between context *Y* and *X*, across conditioning and test days. **(E)** Estimation plot showing the population coupling of deep versus superficial cells in context *X* and *Y*, across conditioning and test days. **(F)** Estimation plot comparing the population coupling of deep versus superficial cells in context *Y* across conditioning days. ***P<0.001, **P<0.01, *P<0.05.

**Figure S8.**
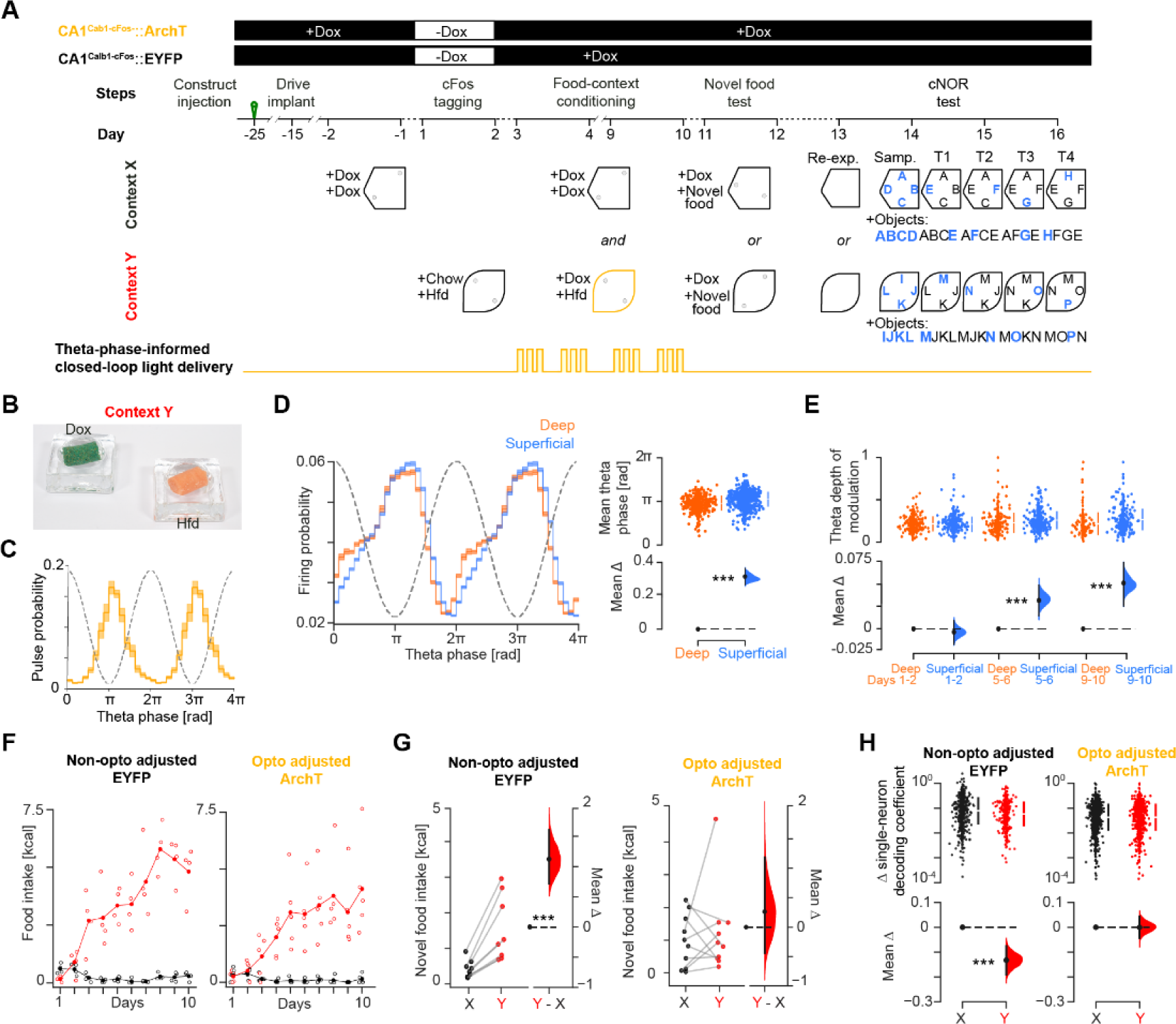
Theta phase-informed closed-loop optogenetic intervention on CA1^Calb1-cFos^::ArchT and CA1^Calb1-cFos^::EYFP mice. **(A)** Experimental layout. We used double-transgenic Calb1-Cre;cFos-tTA mice fed with doxycycline (Dox) containing food pellets in their homecage. Using an intersectional targeting strategy (Figure 4A), we targeted the CA1 of these mice with either the ArchT-EYFP construct or its EYFP-only control; subsequent implantation of a 14-tetrode microdrive including two optic fibers allowed monitoring CA1 neuronal ensemble with theta phase-informed bilateral light delivery. To optogenetically tag CA1 neurons in the Hfd-paired context *Y* with ArchT-EYFP (in optogenetically-adjusted CA1^Calb1-cFos^::ArchT mice) or EYFP-only (in non-optogenetically-adjusted CA1^Calb1-cFos^::EYFP mice), the homecage feeding with Dox pellets was then transiently replaced with regular Chow pellets for the 48-hour period corresponding to days 1 and 2 of Hfd-context *Y* conditioning. To restrict this tagging to context *Y*, mice were not exposed to the other contexts during this off-Dox period. For this reason, mice started their day 1 and 2 conditioning of Chow in context *X* in the two days before, while still on Dox diet (i.e., they were not exposed to context *X* for the two days corresponding to Hfd-context *Y* tagging). CA1 light delivery was then actuated by real time tracking of theta phase while mice continued to explore context *Y* with Hfd from conditioning day 3 onward. **(B)** Picture of the two food containers used to provide mice with a choice between one Dox-diet pellet versus one Hfd pellet in context *Y* (from day 3 conditioning onward). **(C)** Light pulse probability as a function of ongoing theta phase for closed-loop CA1 light delivery during Hfd conditioning in context *Y* (conditioning day 3 onward). **(D)** Left: Firing probability of deep (orange) versus superficial (blue) CA1 *pyramidale* sublayer cells as a function of theta phase. Right: Estimation plot showing the mean theta phase coupling difference between these two subpopulations. Note that superficial cells tend to fire later in the theta cycle. **(E)** Estimation plot showing that the theta modulation of superficial *pyramidale* sublayer cell spiking increases compared to deep *pyramidale* sublayer cells as the Hfd conditioning progresses in context *Y*. **(F)** Food intake over the 10-day conditioning in context *X* (black) versus context *Y* (red) for the non-optogenetically-adjusted CA1^Calb1-cFos^::EYFP control mice (left) and the optogenetically-adjusted CA1^Calb1-cFos^::ArchT mice (right). Each data point represents one mouse. **(G)** Estimation plots showing the effect size for the difference in novel food intake across context *X* and *Y* in the post-conditioning novel food test for the non-optogenetically-adjusted CA1^Calb1-cFos^::EYFP control mice (left) and the optogenetically-adjusted CA1^Calb1-cFos^::ArchT mice (right). Each data point represents one mouse. **(H)** Estimation plots showing the change (update) in the magnitude of single neuron contribution to population novel object-location classification, quantified as the difference of the GLM coefficients in either context (see Figure 1J and methods). ***P<0.001.

